# AGEpy: a Python package for computational biology

**DOI:** 10.1101/450890

**Authors:** Franziska Metge, Robert Sehlke, Jorge Boucas

**Affiliations:** Bioinformatics, Max Planck Institute for Biology of Ageing, Cologne, 50931, Germany; Biological Mechanisms of Ageing, Max Planck Institute for Biology of Ageing, Cologne, 50931, Germany; Cellular Networks and Systems Biology, CECAD, University of Cologne, 50931 Cologne, Germany

## Abstract

**Summary:** AGEpy is a Python package focused on the transformation of interpretable data into biological meaning. It is designed to support high-throughput analysis of pre-processed biological data using either local Python based processing or Python based API calls to local or remote servers. In this application note we describe its different Python modules as well as its command line accessible tools `aDiff`, `abed`, `blasto`, `david`, and `obo2tsv`.

**Availability:** The open source AGEpy Python package is freely available at: https://github.com/mpg-age-bioinformatics/AGEpy.

## 1 Introduction

The generation of meaning from data has become a central topic in biological research. Many tools have therefore emerged for the transformation of raw data into interpretable values in the same way that others have emerged for the generation of biological meaning from such interpretable values. While the firsts are often based on a programmatic use - eg. RNAseq analysis tools like Cuffdiff (Trapnell *et al.*, 2012) or DEseq2 (Love *et al.*, 2014) - the latests tend to be graphical user interface (GUI) based and have more recently evolved to provide access through an application programming interface (API) - eg. KEGG (Kanehisa *et al.*, 2017) and DAVID (Jiao *et al.*, 2012). API access has emerged as a reflection of the growing need to massively process data with tools initially designed for manual analysis through GUIs. APIs offer a way to democratize computational tools with a client side server-less solution.

Here we introduce AGEpy, a Python package for the automation of data analysis at the data-to-meaning interface. It uses standard data science dependencies like pandas (McKinney, 2010), numpy (van der Walt *et al.*, 2011), and matplotlib (Hunter, 2007); bionformatics tools like pybedtools (Dale *et al.*, 2011); and APIs requests to DAVID, blast (Camacho *et al.*, 2009; NCBI Resource Coordinators, 2013), cytoscape (Shannon *et al.*, 2003) and others. Defaults are set for research focused on the biology of ageing using *Caenorhabditis elegans*, *Drosophila melanogaster*, *Mus musculus*, and *Homo sapiens*. AGEpy provides command line executables for downstream analysis of differential gene expression results output by Cuffdiff (Trapnell *et al.*, 2012) - `aDiff` - annotation of bed files - `abed` - online blast queries - `blasto` - DAVID queries - `david` - as well as parsing of gene ontology obo files into tsv - `obo2tsv`.

## 2 Approach

### 2.1 Modules

We have divided AGEpy functions across 13 different modules with most functions making use of standard Python structures as well as data science ones, eg.: numpy arrays and pandas dataframes: `bed.py, blast.py, cytoscape.py, david.py, fasta.py, go.py, gtf.py, homology.py, kegg.py, meme.py, sam.py`. Of notice, `biom.py` (biomart), `cytoscape.py, david.py`, and `kegg.py` make use of the respective service’s APIs. `homology.py` collects homology information from NCBI’s homologene https://www.ncbi.nlm.nih.gov/homologene - to generate homology tables. `plots.py` introduces two visualization plots for enrichment analysis (Figure 1 and 2).

**Figure 1:**
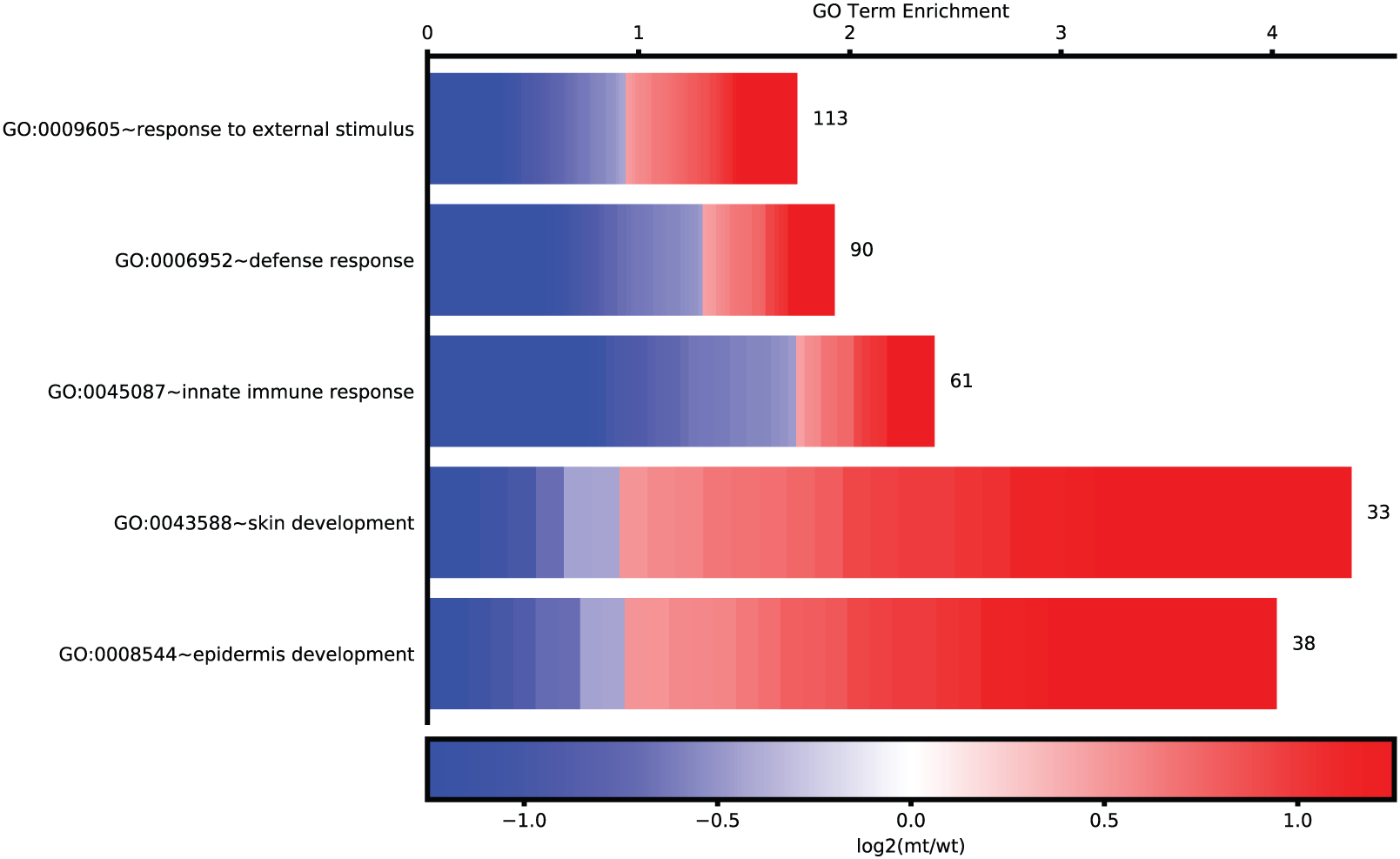
Cellplot. A representation of enriched go terms. Enrichment of top 5 significant terms is plotted on the x-axis. For each term the number of genes is shown as well as the log2(fold change) of each respective gene.

**Figure 2:**
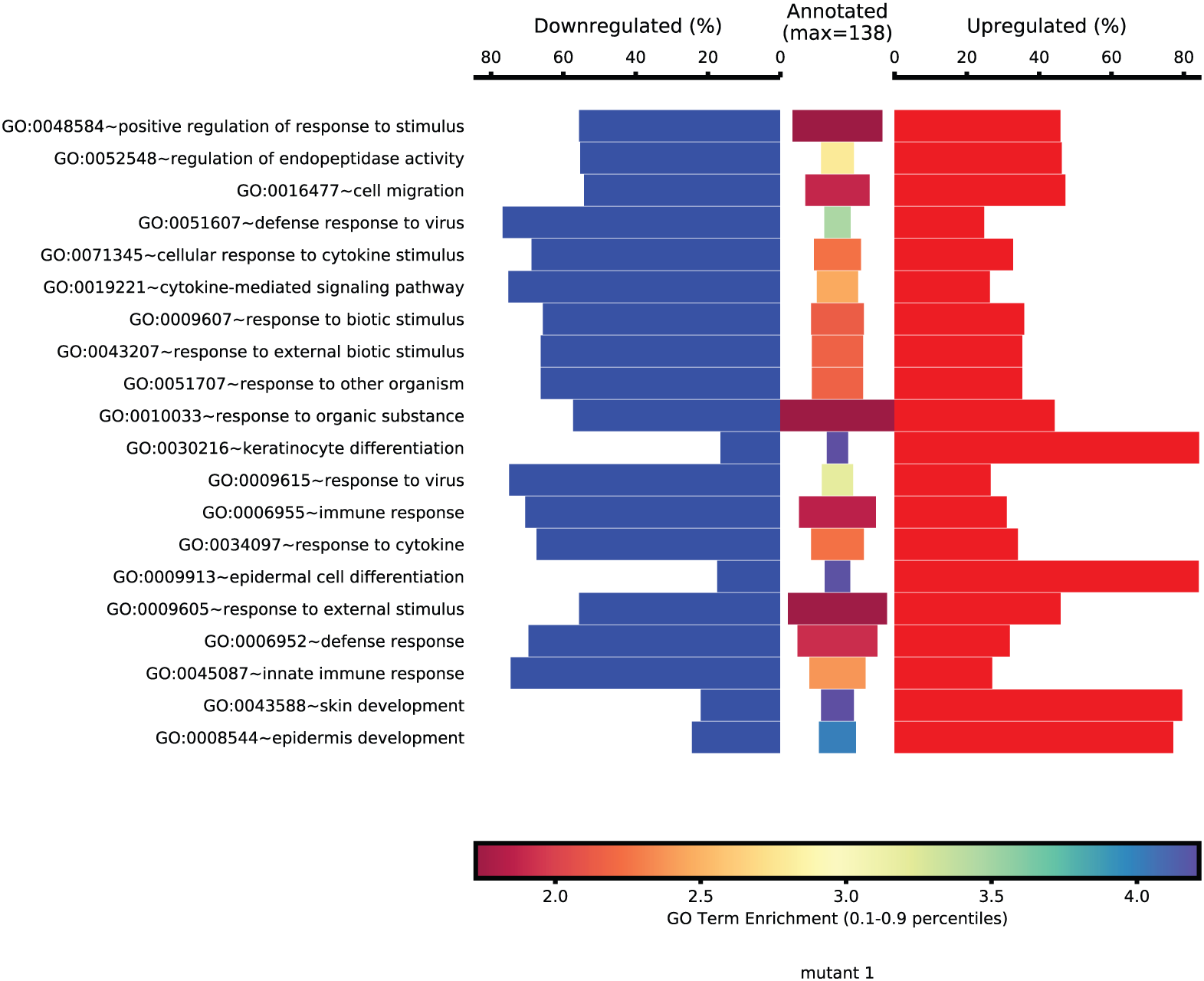
Symplot. A representation of changed genes within an enriched term. Top 20 enriched terms are shown. The size of the central bars is directly proportional to the number of genes in the query belonging to the respective term and the enrichement value is reflected by the color of the bar. Percentage of down and upregulated genes in each term is shown.

### 2.2 Executables

#### 2.2.1 aDiff

`aDiff` is an annotation and mining tool for differential expression results generated with Cuffdiff (Trapnell *et al.*, 2012) - differential gene expression, differential isoforms expression, differential promoter usage, differential splicing, and differential cds. It starts by mapping Cuffdiff’s self-generated artificial gene/transcript ids to gene/transcript ids in a provided *ensembl* reference genome annotation (as provided to Cuffmerge). Using *ensembl* gene ids, it collects biotype and gene ontology information from *ensembl*’s biomart (Smedley *et al.*, 2015) server as well as KEGG annotations through the KEGG API (Kanehisa *et al.*, 2017) for each respective gene. A report of these annotated tables can be obtained in both *tsv* or *excel* format. After discarding non-significant values (as defined in Cuffdiff) it generates one report sheet for each pair-wise comparison. Each respective list of significant genes/transcripts is used to query “The Database for Annotation, Visualization and Integrated Discovery (DAVID)” (Jiao *et al.*, 2012) for enrichment in user defined categories (default: go term bp fat, go term cc fat, got term mf fat, kegg pathway, pfam, prosite, genetic association db disease, omim disease). Protein-protein interaction networks are assembled through the STRING Cytoscape app (Szklarczyk *et al.*, 2017) using cytoscape’s REST API running locally or remotely. An example of an aDiff output from the raw data provided by (Trapnell *et al.*, 2012) can be downloaded from the projects wiki https://github.com/mpg-age-bioinformatics/AGEpy/wiki.

#### 2.2.2 abed

`abed` annotates bed files with overlapped gene names, gene ids, and other defined features (eg. promoter, exon, UTR).

#### 2.2.3 blasto

`blasto` uses the `blast.py` module to query remote blast servers through their respective REST API’s. For complex queries with multifasta files where different arguments are required for the different fasta entries, arguments can be given in the respective sequence names. Tabular results as well as html can be saved locally.

#### 2.2.4 david

`david` uses part of the `david.py` module to query DAVID through its REST API with user provided gene lists in a tabular format. Given a second column with log2(fold change) it will use the CellPlot and SymPlot functions from the `plots.py` module to display the respective plots (Figure1 and 2). As an option DAVID queries can be performed against a user provided list of background genes.

#### 2.2.5 obo2tsv

`obo2tsv` takes annotation files (in obo format) from the gene ontology consortium (Ashburner *et al.*, 2000; The Gene Ontology Consortium, 2017) as input and converts them into a tsv formatted file with parent and child term information included. Given a gene association file, for a selected organism, the generated tabular gene ontology annotation will be merged with the respective organism data. Input can either be local or URLs to the respective files.

## 3 Conclusion

AGEpy provides Python support for computational biology. Use cases for several of its functions have recently been published (Boucas, 2018) and can be further experienced with its provided executables. With most default arguments set for research on the biology of ageing using worm, fly, mouse or human data it can be useful for the interpretation of ageing research data on both small and large scales. AGEpy modules come with extensive documentation (available under the package’s *docs* folder and at http://agepy.readthedocs.io). The open source AGEpy Python package is freely available at https://github.com/mpg-age-bioinformatics/AGEpy.

## Acknowledgements

We acknowledge all the current and past members of the Bioinformatics Core Facility of the Max Planck Institute for Biology of Ageing.

